# SARS-CoV-2 elicits robust adaptive immune responses regardless of disease severity

**DOI:** 10.1101/2020.10.08.331645

**Authors:** Stine SF Nielsen, Line K Vibholm, Ida Monrad, Rikke Olesen, Giacomo S Frattari, Marie H Pahus, Jesper F Højen, Jesper D Gunst, Christian Erikstrup, Andreas Holleufer, Rune Hartmann, Lars Østergaard, Ole S Søgaard, Mariane H Schleimann, Martin Tolstrup

## Abstract

The SARS-CoV-2 pandemic currently prevails worldwide. To understand the immunological signature of SARS-CoV-2 infections and aid the search for treatments and vaccines, comprehensive characterization of adaptive immune responses towards SARS-CoV-2 is needed. We investigated the breadth and potency of antibody-, and T-cell immune responses, in 203 recovered SARS-CoV-2 infected patients who presented with asymptomatic to severe infections. We report very broad serological profiles with cross-reactivity to other human coronaviruses. Further, >99% had SARS-CoV-2 epitope specific antibodies, with SARS-CoV-2 neutralization and spike-ACE2 receptor interaction blocking observed in 95% of individuals. A significant positive correlation between spike-ACE2 blocking antibody titers and neutralization potency was observed. SARS-CoV-2 specific CD8^+^ T-cell responses were clear and quantifiable in 90% of HLA-A2^+^ individuals. The viral surface spike protein was identified as the dominant target for both neutralizing antibodies and CD8^+^ T cell responses. Overall, the majority of patients had robust adaptive immune responses, regardless of disease severity.

**Author summary:** SARS-CoV-2 can cause severe and deadly infections. However, the immunological understanding of this viral infection is limited. Currently, several vaccines are being developed to help limit transmission and prevent the current pandemic. However, basic understanding of the adaptive immune response developed during SARS-CoV-2 infections is needed to inform further vaccine development and to understand the protective properties of the developed immune response. We investigated, the adaptive immune response developed during SARS-CoV-2 infections in recovered patients experiencing a full spectrum of disease severity, from asymptomatic infections to severe cases requiring hospitalization. We used a novel multiplex serological platform, cell-based neutralization assays and dextramer flow cytometry assays to characterize a broad and robust humoral and cellular immune response towards SARS-CoV-2. We found that the vast majority of recovered individuals have clear detectable and functional SARS-CoV-2 spike specific adaptive immune responses, despite diverse disease severities. The detection of both a humoral and cellular functional spike specific immune response in the vast majority of the individuals, irrespective of asymptomatic manifestations, supports vaccine designs currently underway, and encourages further exploration of whether primary infections provide protection to reinfection.

## Introduction

The year of 2020 has been thoroughly marked by the outbreak of severe acute respiratory syndrome Coronavirus 2 (SARS-CoV-2)[1]. Originating in China December 2019, the outbreak was formally declared a pandemic by the WHO in March 2020 [2]. With millions of cases confirmed across 200 countries, the virus has claimed more than 1.4 million lives as of early December 2020 [3]. The SARS-CoV-2 epidemic is an ongoing health crisis, which is extensively affecting almost all aspects of the global human society. An important aspect of SARS-CoV-2 replication is binding and infection of the host cell. The viral spike protein receptor binding domain (RBD) interacts with angiotensin-converting enzyme 2 (ACE2), found on the cell surface, thereby mediating viral infection [4, 5]. Coronavirus Disease 2019 (COVID-19) symptoms manifest primarily as a respiratory disease, with emergent complications of several organs in cases of severe disease [6]. While efforts are converging globally to develop an effective vaccine[7], our broader basic understanding of the adaptive immune response towards SARS-CoV-2 is still limited.

Several studies have described the general adaptive immune responses towards SARS-CoV-2, showing that SARS-CoV-2 specific B and T cells are generated during infections. First immunoglobulin (Ig) M and later IgG SARS-CoV-2 spike specific antibodies are readily detected in COVID-19 patients [8–12]. Evaluations by neutralization assays have confirmed the ability of the generated antibodies to prevent viral infections *in vitro* [13–15]. The limited number of confirmed cases suffering reinfections post recovery [16–19], and high degree of protective immunity against viral re-challenge shown *in vivo* in macaque challenge studies [20], suggest that the immunological response developed during primary infections provide at least some protection against reinfection. Additionally, SARS-CoV-2 specific T-cell activation has also been documented in a range of studies [21–23]. However, most studies have been limited to specific disease severity populations, and small or none RT-PCR verified cohorts.

Currently, in depth characterization of the adaptive immune response to SARS-CoV-2 in a large cohort representing the full disease spectrum, as well as the development of functional, and easily scalable, serological assays, are needed to guide and support rapid further vaccine development. Here, we have delineated the humoral and cellular immune responses in 203, RT-PCR verified, recovered SARS-CoV-2 patients. We evaluated the quantity and potency of antibodies in each individual towards several different coronaviruses and antigens, using both a SARS-CoV-2 spike pseudovirus neutralization assay and a novel Mesoscale Diagnostics (MSD) multiplex platform [24]. We further quantified the breadth and magnitude of single-epitope SARS-CoV-2 specific CD8^+^ T cells, using dextramer flow cytometry. Thus, we report an extensive panel of adaptive immune parameters in the context of disease severity, to provide an outline of the general broad and functional SARS-CoV-2 specific adaptive immune response observed across the full COVID-19 disease spectrum.

## Results

### Patient enrollment

We studied the adaptive immune response towards SARS-CoV-2 among 203 patients who had recovered from COVID-19. We have recently described the cohorts clinical characteristics thoroughly [25] a basic overview of which is shown in table 1. The median age of individuals was 47 years (range: 21 – 79), and 45% were female. The cohort was divided into three COVID-19 disease severity groups. 1: Home/outpatients with no limitation of daily activities (8%), 2: Home/outpatients with a limitation of daily activities (75%), and 3: Hospitalized patients (17%). The median duration of COVID-19 symptoms was 13 days (range: 0 – 68). Enrollment occurred at least 14 days after the end of COVID-19 related symptoms, with a median of 31 (range: 14 – 61) days from time of recovery to study enrollment. To allow comparison of immunological outcomes from SARS-CoV-2 infection recovered patients, samples from 10 healthy individuals enrolled in a study conducted prior to the current COVID-19 pandemic were included as controls [26].

**Table 1:** Demographics and Clinical Characteristics at Baseline, Cohort characteristics. All individuals were assigned a COVID-19 severity group depending on their course of disease. Group 1 consisted of asymptomatic individuals with no limitations in their daily activities. Group 2 of moderately sick, able to recover at home. Finally, group 3 comprises all hospitalized individuals, including those with/without oxygen requirement and/or ICU admission.

| **Table 1: Demographics and Clinical Characteristics at Baseline** |  |  |
| --- | --- | --- |
| **Characteristics** | **n=203** |  |
| Age, years, median (range) | 47 | (21-79) |
| Female sex, no (%) | 92 | (45) |
| HLA-A2<sup>+</sup>, no (%) | 113 | (56) |
| COVID-19 disease severity, no (%) |  |  |
| 1. Home/outpatient, no limitation of daily activities (asymptomatic/mild) | 17 | (8) |
| 2. Home/outpatient, limitation of daily activities (moderate) | 152 | (75) |
| 3. Hospitalized (severe) | 34 | (17) |
| Duration of COVID-19 symptoms, days, median (range) | 13 | (0-68) |
| Time from recovery to inclusion, day, median (range) | 31 | (14-61) |

### Human coronavirus serology

First, we analyzed the presence of IgG antibodies towards multiple human coronaviruses in serum, using the multiplex MSD platform. Compared to controls, we found significantly elevated levels of IgG antibodies in spike RBD, spike N-terminal domain (NTD), and the nucleocapsid (p<0.0001, Fig 1A). Furthermore, IgG antibodies from SARS-CoV-2 infected individuals exhibited strongly increased reactivity towards spike protein from other human beta coronaviruses: SARS-CoV-1 and Middle East respiratory syndrome (MERS), as compared to the controls. Further, increased IgG levels towards the seasonal beta coronavirus strains: HKU1 and OC43, compared to IgG from the control group were also observed (p<0.0001, Fig 1B). No difference was detected in IgG levels to the negative bovine serum albumin (BSA) control between SARS-CoV-2 patients and controls. Importantly, 202 out of the 203 individuals analyzed here, developed detectable antibodies, otherwise absent in the historical controls, against both full-length SARS-CoV-2 Spike and RBD antigens, during SARS-CoV-2 infections. Likewise, robust production of IgA antibodies was also observed for nearly all infected individuals, with SARS-CoV-2 spike specific IgA levels being significantly elevated compared to controls in 201 of the 203 individuals (Fig 1C). Additionally, SARS-CoV-2 IgG levels towards both spike and nucleocapsid antigens, correlated positively with the disease severity. (Fig 1D+E). Overall, we conclude that more than 99% of the SARS-CoV-2 infected individuals in this cohort had readily detectable antibodies to SARS-CoV-2 spike antigen, and that broad IgG immunological recognition of SARS-CoV-2 with cross-reactivity to several different coronavirus develops during COVID-19. Additionally, the magnitude of spike-targeting antibodies increases with disease severity.

**Figure 1:**
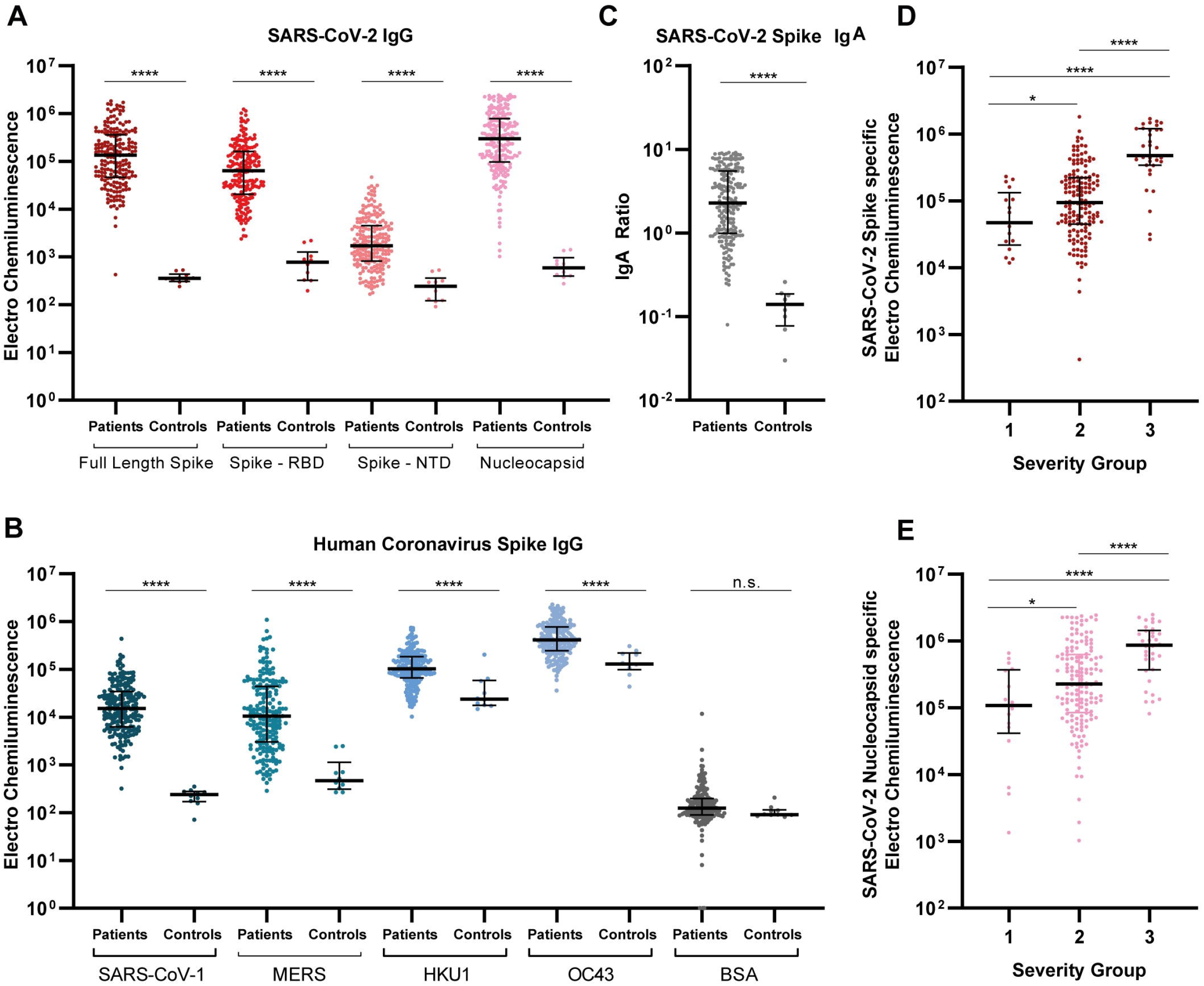
Extensive IgG and IgA presence with multiple SARS-CoV-2 antigens. **A+B)** Serum IgG levels for all individuals and 10 pre-pandemic healthy controls. IgG was detected against SARS-CoV-2 Spike, RBD (receptor binding domain), NTD (N-terminal domain), nucleocapsid and non-SARS-CoV-2 spike proteins of other corona virus. Data are blank-corrected electro chemiluminescent signal measured by MSD multiplex serology assays. **C)** Serum IgA levels for all individuals and eight pre-pandemic healthy controls, measured by ELISA. IgA is shown as a ratio against a standard calibrator. **D+E)** Distribution of IgG volumes between each disease severity group, for both SARS-CoV-2 spike (D) and nucleocapsid (E). Data are blank-corrected electro chemiluminescent signal measured by MSD multiplex serology assays. Scatter plots with individual data points are shown with median (wide line) and interquartile range (narrow lines). Statistical comparison between groups were done by Mann-Whitney U test. n.s = not significant, * =p<0.05, **** = p<0.0001, n = 203.

### SARS-CoV-2 pseudovirus neutralization

Next, we investigated the functional neutralization capacity of total plasma antibodies *in vitro,* using VSV pseudotyped virus expressing SARS-CoV-2 spike protein. Antibody neutralizing potency was evaluated by serial dilutions of plasma, yielding infectivity titration curves for each of the SARS-CoV-2 infected individuals and the controls (Fig 2A). We found that 95.5% of the individuals (193 of 202) were able to neutralize SARS-CoV-2 spike pseudoviruses and provided 100% inhibition at the lowest (1:25) plasma dilution. IC50 values were extrapolated from the neutralization curves, and assigned to each individual as a measure of antibody neutralization potency. Serum from the remaining nine individuals (4.5%) were unable to fully neutralize viral infection, producing neutralization curves comparable to that of the uninfected controls. No legitimate IC50 value could be calculated for these individual, and consequently they were excluded from further analyses using this parameter. Collectively, the IC50 values of all 193 neutralizing individuals span evenly across four orders of magnitude (Fig 2B). In concurrence with the analysis in Fig 1D+E, we observed lower IC50 values among individuals experiencing mild symptoms compared to those with moderate (p<0.001) or severe COVID-19 (p<0.0001) (Fig 2C). We conclude that in this large cohort, with considerable diversity in disease severity, the vast majority (>95 %) of SARS-CoV-2 infections lead to the production of effective neutralizing antibodies, and that neutralization potency increases with disease severity.

**Figure 2:**
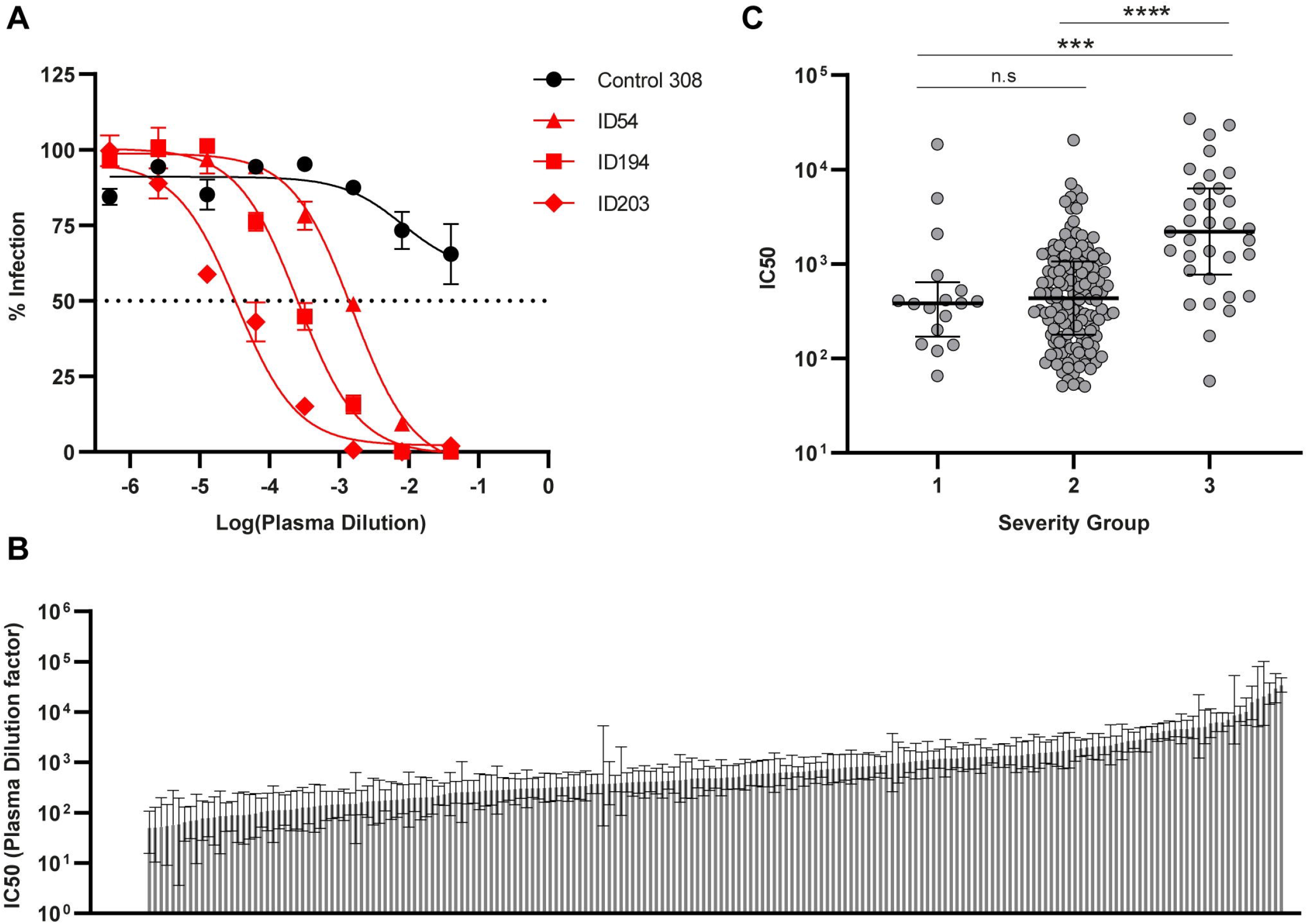
SARS-CoV-2 neutralization capacity correlates with disease severity. **A)** Representative neutralization curves for control ID308, and individuals ID54, ID194, and ID203, quantified as eGFP^+^ cells by flow cytometry. Control plasma was unable to neutralize below a 50% infection rate, where SARS-CoV-2 recovered patients accomplish 100% neutralization at the lowest plasma dilution. X-axis shows the log10 transformed patient plasma dilution, from 1:25 – 1:1,953,125. Error bars represent mean and s.e.m. of duplicate determinations. Three-parameter non-linear fit is plotted. **B)** IC50 values calculated from neutralization curves, graphed from lowest (left) – highest (right) within the cohort. Error bars show 95% confidence interval. Nine individuals unable to neutralize 100% are represented with the value zero on the y-axis far left, n = 202. **C)** Distribution of IC50 values between disease severities. Scatter plot with individual data points shown with median (wide line) and interquartile range (narrow lines). Statistical comparison were by Mann-Whitney U test. *** = p < 0.001, **** = p < 0.0001, n = 193.

### Antibodies efficiently block ACE2 receptor binding

We continued the characterization of SARS-CoV-2 antibody functionality, using an MSD SARS-CoV Spike – ACE2 competition assay (Fig 3A). This allowed us to measure the quantity of antibodies able to block the interaction between the ACE2 receptor and SARS-CoV-2 full-length spike protein, SARS-CoV-2 RBD, and SARS-CoV-1 spike protein. Many of the recovered individuals reached the assay’s upper limit of quantification, and a clear increase in the quantities of serum ACE2 blocking antibodies was observed for all three antigens compared to historic controls (p≤0.0001) (Fig 3B). The levels of antibodies blocking SARS-CoV-2 Spike – ACE2 receptor interaction was increased in >99% of the individuals (202 of 203) compared to uninfected controls. The individual antibody concentrations also correlated to the time from disease recovery to inclusion (S1 Fig 1). Nevertheless, we found that those experiencing severe COVID-19 had significantly greater levels of SARS-CoV-2 spike specific ACE2 blocking antibodies, compared to individuals with mild to moderate disease (p<0.0001, Fig 3C). Both the pseudovirus cell-based neutralization assay and the SARS-CoV Spike – ACE2 competition assay investigate the presence of functional antibodies towards SARS-CoV-2. We identified a highly significant correlation between the IC50 values from the pseudovirus neutralization assay and the concentration of SARS-CoV-2 spike specific antibodies capable of blocking ACE2 receptor interaction (p>0.0001 Fig 3D). In conclusion, we observed that nearly all individuals produce antibodies that target the spike protein-ACE2-receptor interaction and that the level of these antibodies was increased with severe disease. Further, the virus neutralization capacity increased in conjunction with the amount of functional ACE2 blocking antibody present in serum.

**Figure 3:**
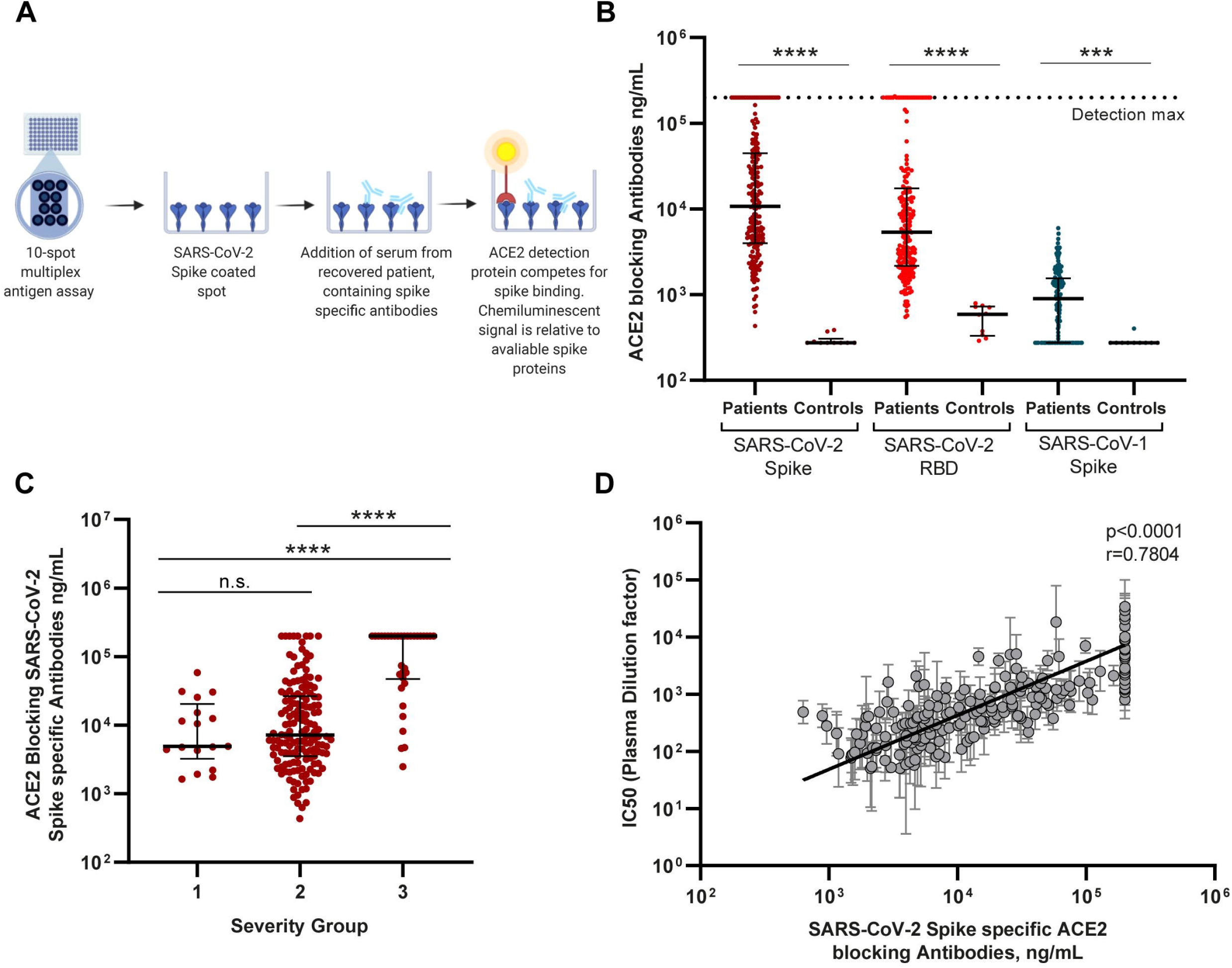
SARS-CoV-2 antibody quantification by ACE2 competition assay. **A)** Schematic drawing of the MSD ACE2 competition assay. Spike-specific serum antibodies bind to their respective epitopes, blocking SULFO-Tag conjugated ACE2. Antibody concentration in ng/ml is calculated based on internal standard antibody blocking ACE2 binding. **B)** Serum ACE2 blocking antibody levels detected against SARS-CoV-2 Spike and RBD, and SARS-CoV-1 spike proteins. Scatter plot with individual data points shown with median (wide line) and interquartile range (narrow lines). Statistical comparison by Mann-Whitney U test. *** = p < 0.001, **** = p<0.0001, n = 203. **C)** Distribution of SARS-CoV-2 spike specific ACE2 blocking antibodies between disease severity groups. Scatter plot with individual data points shown with median (wide line) and interquartile range (narrow lines). Statistical comparison by Mann-Whitney U test. *** = p < 0.001 **** = p < 0.0001, n = 203. **D)** Correlation analysis of pseudotype virus neutralization IC50 values and the quantity of SARS-CoV-2 spike specific ACE2 blocking antibodies. Correlation by Spearman’s rank coefficient, p < 0.0001. n = 193.

### Collected serological analysis

Next, we constructed a heatmap compiling all humoral immunological data, to gain a cohort wide perspective of the overall antibody response developed during SARS-CoV-2 infection. We ranked individuals according to their antibody response potency from the pseudovirus neutralization assay (IC50 value), displaying their respective immunological variables underneath (Fig 4). We observed, that the neutralization capacity was clearly linked to the overall antibody levels present in the patients. Interestingly, it was further evident, that the best (top 10%) neutralizers of the cohort displayed a corresponding increase in the overall breadth of their antibody response, towards all the investigated coronavirus antigens. Importantly, strong pseudovirus neutralization profiles were almost exclusively seen in individuals with antibodies that potently block spike-ACE2 receptor interaction. We therefore conclude that the best neutralizers exhibit a broader variety of cross-reactive antibodies and have greater levels of spike binding and receptor-blocking antibodies.

**Figure 4:**
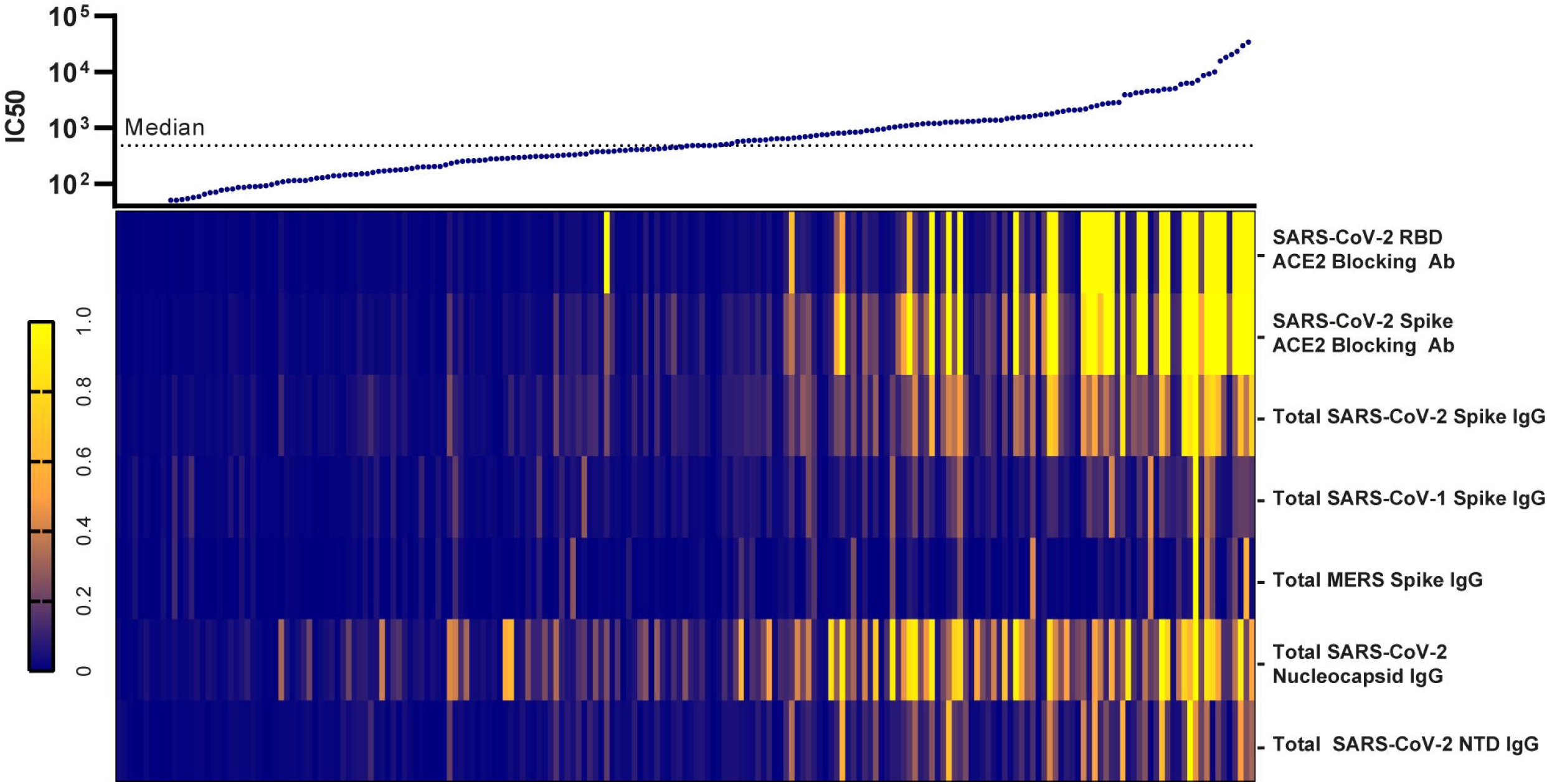
The breadth of immunological response shifts in conjunction with neutralization capacity. Presentation of all IC50 values listed from lowest (left) to highest (right) with a heatmap representing the individuals corresponding relative IgG levels and ACE2 blocking antibody quantities collected through MSD analysis. The normalization of variables within each measured immunological parameter was performed by assigning the highest values to one (bright yellow) and the lowest value to zero (dark blue). n=202.

### Epitope specific CD8+ T cell-responses

We then went on to explore the epitope specific T-cell responses in SARS-CoV-2 recovered individuals. We analyzed the reactivity of CD8^+^ T cells from 106 HLA-A2^+^ individuals in the cohort for their specificity to nine different SARS-CoV-2 epitopes using dextramer staining flow cytometry (Fig 5A). Overall, Membrane_61-70_ (M) (epitope 1), Nucleocapsid_222-230_ (N) (epitope 3), and Spike_269-277_ (S) (epitope 6) were the most commonly recognized epitopes with positive responses detected in 17%, 25% and 81% of individuals, respectively (Fig 5B). Interestingly, these three epitopes originate from three separate SARS-CoV-2 proteins (Fig 5A). The frequency of SARS-CoV-2 specific CD8^+^ T-cells was similar across all nine HLA-A2+ epitopes tested, with the highest individual responses observed for N_222-230_ and S_269-277_ (epitopes 3 and 6) (Fig 5C). Only 10% of the HLA-A2^+^ individuals (11 of 106) had no detectable response to any of the epitopes tested, while the remaining 90% responded to at least one, and up to seven, of the analyzed epitopes. (Fig 5D). We compared the cumulative frequency of SARS-CoV-2 specific CD8^+^ T cells across the disease severity groups and observed no significant difference (Fig 5E). However, we did observe significant albeit weak correlations between the cumulative frequency of SARS-CoV-2 specific CD8^+^ T-cells and the majority of the serological immunological parameters analyzed, including pseudovirus neutralization IC50 values as well as SARS-CoV-2 specific antibody production and ACE2 blocking ability, as outlined with correlation coefficients in table 2. The 11 individuals with no detectable CD8^+^ T-cell responses were evenly distributed among the disease severity groups and displayed varying antibody neutralization capacity (S1 Fig 3). Based on this we were only able to identify two individuals with both no detectable neutralizing antibodies and no detectable CD8^+^ T-cell responses. Thus, we conclude that 90% of SARS-CoV-2 infected individuals mount a detectable CD8^+^ T cell response, towards the nine epitopes tested, irrespectively of disease severity. We further conclude that the broadest targeted epitope in this cohort is located in the spike protein. Lastly, there is an overall weak but statistically significant correlation of antibody responses and CD8^+^ T-cell responses.

**Figure 5:**
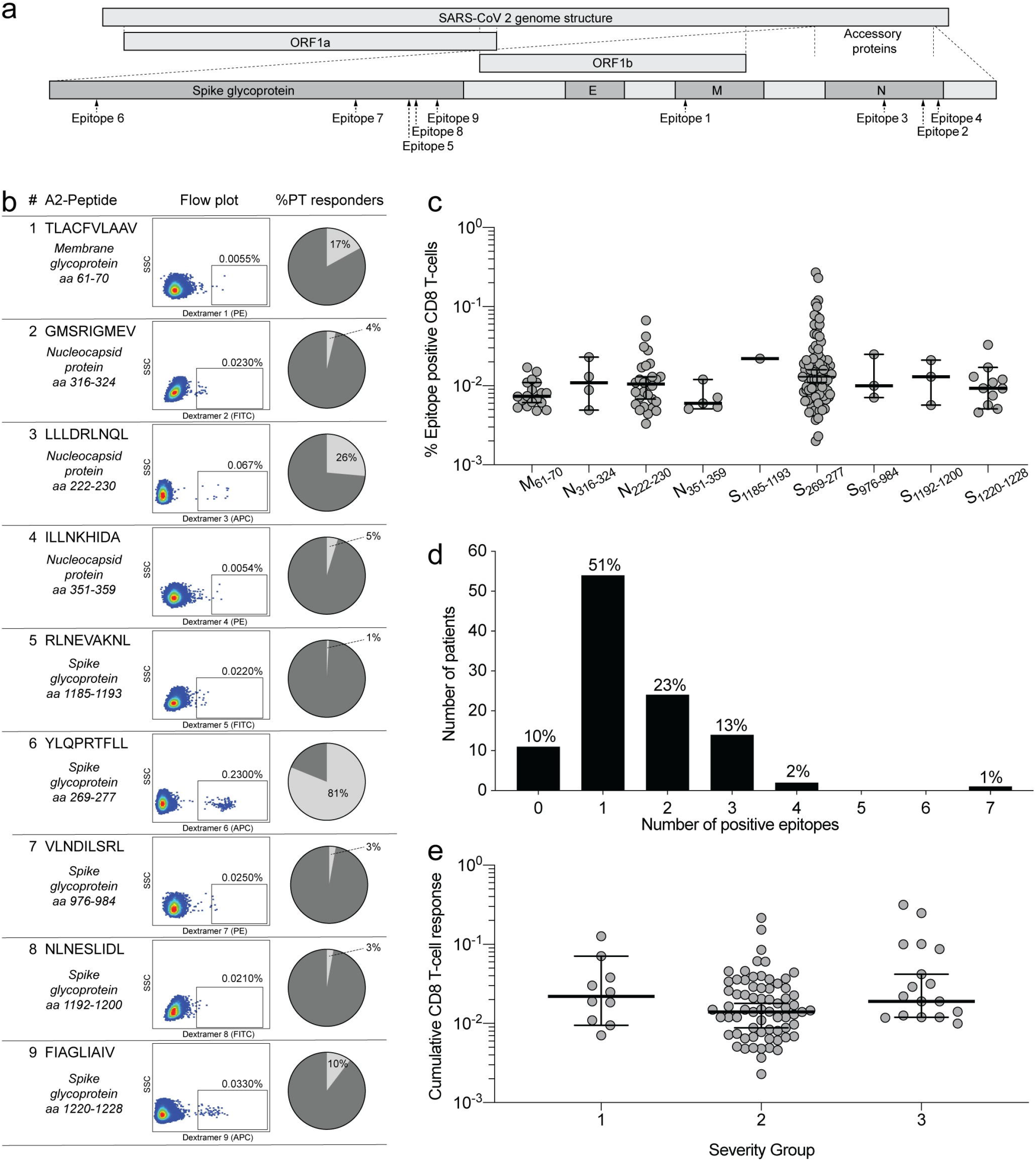
Characterization of CD8^+^ T-cell responses towards SARS-CoV-2 in HLA-A2^+^ individuals. **A)** Overview of HLA-A2^+^ epitope location within the SARS-CoV-2 proteins. **B).** Epitope sequence and individual dextramer signal gating strategy on CD8^+^ T cells, with the percentage of recognition within the cohort shown for each. Full gating strategy is displayed in S1 Fig 2. **C)** The frequency of SARS-CoV-2 responsive CD8^+^ T-cells for each epitope. Scatter plot with individual data points shown with median (wide line) and interquartile range (narrow lines). n = 106 **D)** Breadth of CD8^+^ T-cell responses shown as the cumulative number of CD8^+^ T-cell epitopes targeted by patients. Percentage equivalents of patient numbers are indicated on top of the bars for each cumulative group. n = 106 **E)** Distribution of the cumulative CD8^+^ T-cell responses in HLA-A2^+^ individuals, between the disease severity groups. Error bars show median (wide line) and interquartile range (narrow lines). n=106. 10% of individuals had no detectable CD8^+^ T-cell epitope response, and are not shown on the graph but were included in statistical tests. Statistical comparison by Mann-Whitney U test. n.s. = p > 0.05.

**Table 2:**
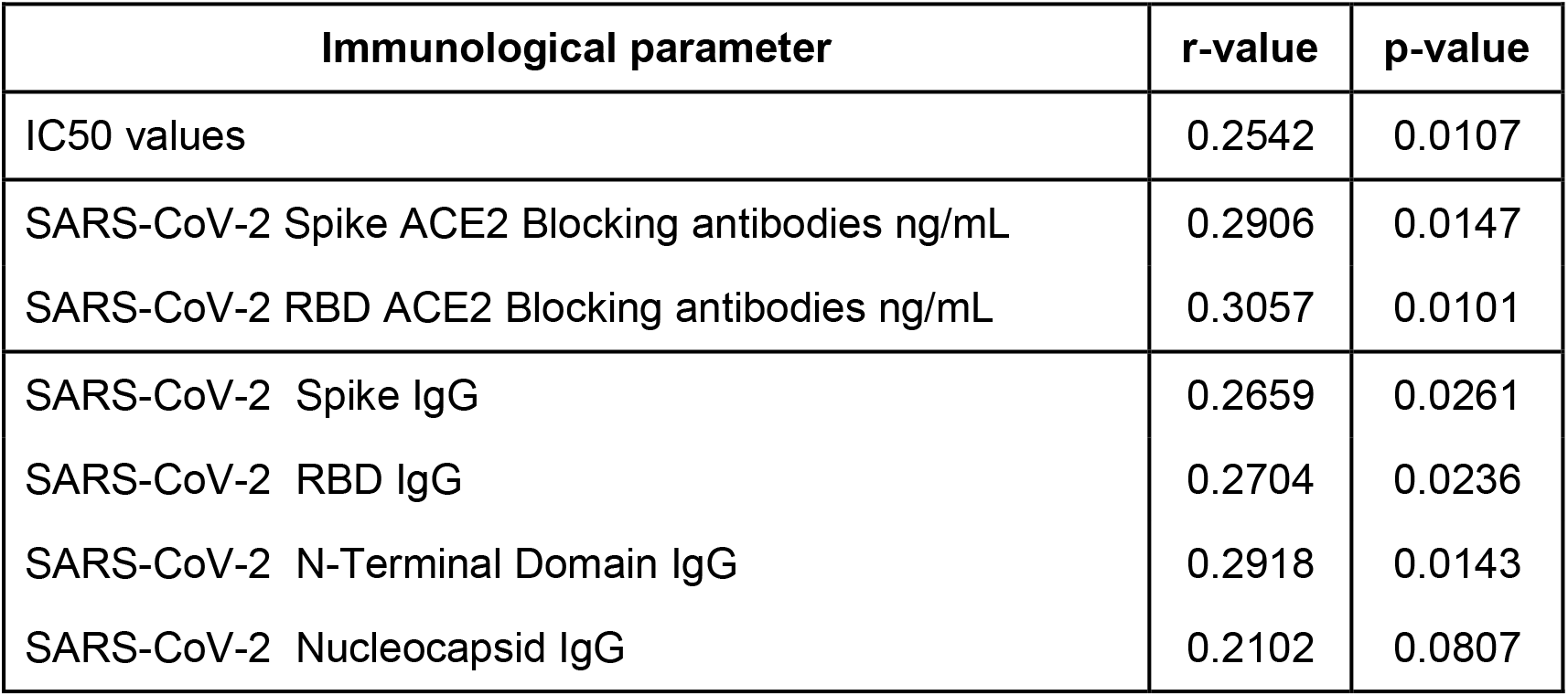
Correlations to cumulative epitope specific CD8+ T-cell responses, Cumulative CD8^+^ T-cell responses in correlation to serology. Spearman’s rank coefficient correlations displaying the relationship between the overall magnitude of CD8^+^ T-cell responses to SARS-CoV-2 epitopes, and antibody neutralization, quantity and ACE2 blocking capacity, for all SARS-CoV-2 antigens investigated.

| **Table 2: Correlations to cumulative epitope specific CD8+ T-cell responses** |  |  |
| --- | --- | --- |
| **Immunological parameter** | **r-value** | **p-value** |
| IC50 values | 0.2542 | 0.0107 |
| SARS-CoV-2 Spike ACE2 Blocking antibodies ng/mL | 0.2906 | 0.0147 |
| SARS-CoV-2 RBD ACE2 Blocking antibodies ng/mL | 0.3057 | 0.0101 |
| SARS-CoV-2 Spike IgG | 0.2659 | 0.0261 |
| SARS-CoV-2 RBD IgG | 0.2704 | 0.0236 |
| SARS-CoV-2 N-Terminal Domain IgG | 0.2918 | 0.0143 |
| SARS-CoV-2 Nucleocapsid IgG | 0.2102 | 0.0807 |

## Discussion

We aimed to characterize the cellular and humoral adaptive immune response in a large cohort of RT-PCR verified SARS-CoV-2 recovered patients, spanning a full spectrum of COVID-19 severity. Overall, our results show that the majority of patients developed a robust and broad both humoral and cellular immune response to SARS-CoV-2. However, our data may also help explain that some rare individuals have no detectable immunological memory to SARS-CoV-2, and will therefore be at risk of re-infection as it has been reported in a few case reports[16–19].

We were able to detect SARS-CoV-2 specific antibodies in all but one of the 203 individuals investigated, irrespectively of their disease severity and duration of symptoms. Antibody specificity was distributed across several SARS-CoV-2 antigens, and with cross coronavirus serological activity observed against SARS-CoV-1, MERS, HKU1, and OC43 human coronaviruses. We assume this reflects cross-reactivity of the antibodies generated against SARS-CoV-2 for two reasons: First, due to the clear significant difference to the pre-pandemic controls, and secondly because no cases of SARS-CoV-1 or MERS have been documented in Denmark. We interpret this as an indication of extensive and broad immune recognition development in COVID-19 patients. Similar to previous studies [14, 27], we confirmed the functionally neutralizing and ACE2 blocking capabilities of the SARS-CoV-2 spike and RBD specific antibodies. Noticeably, this infers the development of a robust humoral immune response within the vast majority of the COVID-19 recovered population. Furthermore, nearly all individuals also have SARS-CoV-2 specific IgA responses, clearly indicating a functional rigorous class switching and maturation. This presence of IgA is crucial for the immunological protection at mucosal barriers, and hence protection against future SARS-CoV-2 exposures.

All serological and functional data collected show that both antibody levels and neutralization potency correlate significantly with the disease severity. This indicates that severe disease manifestation is not caused by a lack of adaptive immunity, which is in line with previous reports [28, 29]. Hence, we suggest that the prolonged disease course, and consequent larger exposure to virus experienced in hospitalized patients, may provide a timeframe in which enhanced antibody affinity maturation takes place, compared to shorter course mild infections.

Studies are conflicted on the degree to which cross-reactive immunity between different coronavirus develop during SARS-CoV-2 infections [11, 13, 14, 28, 30–33]. The considerable diversity of antigen recognition independent of COVID-19 severity shown here, demonstrates that at least some immunological cross-recognition of several different coronavirus is developed during SARS-CoV-2 infections. This is in line with data on cross-reactivity in CD4^+^ T-cell epitopes between seasonal coronaviruses and SARS-CoV-2 [34]. The cross-reactivity observed between SARS-CoV-2, SARS-CoV-1 and MERS, may be due to conserved epitopes between these viruses, as prior infections with SARS-CoV-1 or MERS within our cohort are highly unlikely.

Such potential cross-reactivity could arise through either newly generated SARS-CoV-2 specific antibodies reacting with conserved epitopes, or by reactivation of memory cells originally generated against seasonal coronaviruses, followed by affinity maturation. Importantly, the multiplex serological analyses we performed do not provide insight into the SARS-CoV-2 antibody response on a monoclonal antibody level. Here, further studies are needed to determine possible protective and cross-reactive properties of single-antibody specificities.

We functionally verified the antibody responses in all individuals, using two separate assays. The cell-based pseudovirus neutralization assays are at present the standard method for determining SARS-CoV-2 neutralizing antibody potency. We additionally used the MSD novel coronavirus multiplex assay, recently reported by Johnson *et al* [24] to determine the ACE2 blocking capability of individual serum antibodies. The significant correlation between the two assay readouts identifies the plate format ACE2 competition assay as a powerful, high-throughput, screening tool, with applications in both SARS-CoV-2 therapeutic neutralizing antibody development, and assessments of functional protective antibody induction in vaccine studies. An immense global effort is currently undertaken to develop effective vaccines against SARS-CoV-2, the majority of which are centered on inducing spike or RBD antigen specific immunity [35]. Here we demonstrate that SARS-CoV-2 spike specific, ACE2 blocking antibodies are found in the majority of infected individuals. Their extensive induction, even in short-term, asymptomatic infections, align with current vaccines designs inducing protective immunity based on spike antigens [36, 37]. Nevertheless, the protective effect of antibodies elicited during natural infections, remains to be determined.

We further report, with single-epitope resolution, a SARS-CoV-2 specific CD8^+^ T-cell response in 90% of the HLA-A2^+^ individuals analyzed. This corresponds well with other studies reporting CD8^+^ T cell activation in 70%–100% of recovered patients using full protein overlapping peptide stimulation [21, 38]. The location of the top three immunogenic epitopes within separate proteins in the viral proteome additionally reinforces our conclusion that a broad immune response is generated towards SARS-CoV-2 in the general infected population. T-cell immunity to SARS-CoV-1 is known to persist for up to six years, in contrast to B-cell immunity [39, 40], underlining the importance of developing protective cell based immunity to SARS-CoV-2 if long term viral protection is to be reached. As an important point, the most broadly recognized CD8^+^ T-cell epitope (S_269-277_) within our cohort (responses detected in 81% of HLA-A2^+^ individuals) is located in the spike antigen. Thus, such epitope specificity can clearly be used to evaluate CD8^+^ T-cell immunity in spike focused vaccine developments currently underway.

Surprisingly, we found that the cumulative CD8^+^ T-cell response, across all epitopes, did not vary by disease severity in contrast to what we, and others [41], observed with antibody levels. While the limited coverage of epitopes investigated here may influence this observation, recent evidence suggests that persistent viral replication in otherwise recovered patients may be linked to CD8^+^ T-cell response magnitude [25]. Despite the different observations with regard to immune responses and disease severity, we found overall significant relationships between humoral and T-cell based immunity, but all of modest strength. A possible explanation could be a synchronized waning of the magnitude of response for both immune parameters during the time from recovery to study enrollment.

Of note, the use of dextramer staining is limited by inclusion of selected epitopes only, and conclusions are consequently limited to the relative low epitope coverage. However, the advantages of the dextramer technology are superior sensitivity and a high degree of specificity. In the light of the relative low proteome coverage, the fact that only 10% of the investigated individuals did not have a detectable CD8^+^ T-cell response clearly indicate a strong cytotoxic T-cell component in the immune response towards SARS-CoV-2. Furthermore, as our observations of breadth and magnitude in relation to the distribution of distinct SARS-CoV-2 antigens are similar to others [38, 41] we conclude that the panel of dextramers applied here provide a new and sensitive representation of the general CD8^+^ T-cell response to SARS-CoV-2 that will be an important tool in assessing long-term immunity following primary infection or vaccination.

In conclusion, we observed that disease severity is closely related to the potency and breadth of the antibody response towards SARS-CoV-2. Furthermore, we identified the SARS-CoV-2 spike protein as a target of adaptive immunity in >99% of the cohort, irrespective of COVID-19 symptom manifestation. Only two individuals (<2%) had neither antibodies with virus neutralization capacity, nor detectable CD8^+^ T-cell responses. Hence, we conclude that regardless of COVID-19 severity, a robust adaptive immune response towards SARS-CoV-2 is elicited during primary infections.

## Materials and Methods

### Study design and sample collection

Samples were collected from a cohort of 203 individuals who had recovered from COVID-19. Participants were enrolled at Department of Infectious Diseases at Aarhus University Hospital, Denmark from April 3^rd^ to May 29^th^ 2020. Inclusion criteria were as follows; 1) Age above 18 years; 2) PCR verified SARS-CoV-2 within the preceding 12 weeks; 3) Full recovery from acute COVID-19 illness; 4) Able to give informed consent. Exclusion criteria were; 1) Ongoing febrile illness; 2) Immunosuppressive treatment and/or known immunodeficiency; 3) Pregnancy. Samples were collected at least 14 days after recovery and a maximum of 12 weeks after SARS-CoV-2 PCR-verified diagnosis. One patient ID116 only had serum collected, and thus is absent from IC50 and T-cell analyses.

Individuals were allocated to three groups according to the severity of COVID-19 illness, based on the criteria: 1) Home/outpatient, not experiencing any limitations in daily activities; 2) Home/outpatient, certain limitations in daily activity level (fever, bedridden during illness); 3) All hospitalized patients, regardless of need for supplemental oxygen treatment, or ICU admission with/without mechanical ventilation. Additional data regarding demographic and clinical characteristics of this cohort has been reported elsewhere [25].

### Serology

IgG antibodies were measured in serum samples using the MSD Coronavirus Plate 1 Cat. No. N05357A-1, MesoScale Discovery, Rockville, Maryland), a solid phase multiplex immunoassay, with 10 pre-coated antigen spots in a 96-well format, with an electro-chemiluminescence based detection system. The SARS-CoV-2 related antigens spotted were CoV-2 Spike, CoV-2 RBD, CoV-2 NTD, and CoV-2 nucleocapsid. The remaining spots comprised antigens from other respiratory pathogens: Spike protein from SARS-CoV-1, MERS coronavirus, and two seasonal coronaviruses OC43, HKU1. BSA served as negative control, as previously described [24]. Unspecific antibody binding was blocked using MSD Blocker A (Cat. No. R93AA-1). COVID-19 patient serum samples and control samples were diluted 1:4630 in MSD Diluent 100 (Cat. No. R50AA-3). After sample incubation, bound IgG was detected by incubation with MSD SULFO-TAG Anti-Human IgG Antibody and subsequently measured on a MESO QuickPlex SQ 120 Reader (Cat. No. AI0AA-0) after addition of GOLD Read Buffer B (Cat. No. R60AM-2).

### ACE2 Competition Assay

Spike and RBD targeting antibodies with the ability to compete with ACE2 binding were measured using the MSD Coronavirus Plate 1. COVID-19 blocking antibody calibrator and 1:10 diluted patient and control serum samples were incubated after plate blocking. SULFO-Tag conjugated ACE2 was added before washing, allowing ACE2 to compete with antibody binding to spike and RBD antigens immobilized on the plate. Bound ACE2 was detected as described for the serology assay above, and antibody concentrations were subsequently calculated using the MSD Discovery Workbench software.

### ELISA

IgA antibodies were measured using the Anti-SARS-CoV-2 IgA ELISA from Euroimmun (Euroimmun Medizinische Labordiagnostika AG, Lübeck, Germany, Cat. No. El 2606-9601 A), according to manufacturer’s instructions. In brief, antibodies in serum samples diluted 1:200 were captured by recombinant S1 domain of SARS-CoV-2 spike protein immobilized in microplate wells. IgA type antibodies were detected by incubation with peroxidase labelled anti-human IgA followed by a chromogen solution, resulting in color development in positive wells. Signal was read at 450 nm with reference measurements at 650 nm, which were used for background signal corrections. Results were analyzed relative to the ELISA kit calibrator, as a ratio between sample absorbance and calibrator absorbance.

### Cells and plasmids

All cell lines were incubated at 37 °C and 5 % CO2 in a humidified atmosphere. BHK-G43, previously described *[42, 43],* were cultured in Dulbecco’s modified eagle’s medium (DMEM), containing 5 % Fetal Bovine Serum (FBS) and 50 U/mL Penicillin G/Streptomycin (P/S), where Zeocin (100μg/ml) and Hygromycin (50μg/ml) were added at every fourth passage. Induction of VSV-G glycoprotein was performed with 10^−8^M mifepristone. HEK293T cells were cultured in DMEM, containing 10% FBS and 50 U/mL P/S. Vero76 cmyc hTMPRSS2 [4] cells were cultured in DMEM supplemented with 10% FBS, 50 U/mL P/S, and 10 μg/mL Blasticidin.

The construction of pCG1-SARS-2-Spike has been previously described [4, 44]. Briefly, SARS-2-S (NCBI Ref.Seq: YP_009724390.1) coding sequence was PCR-amplified and cloned into the pCG1 expression vector via BamHI and XbaI restriction sites.

### Virus production

For generation of VSV*ΔG(luc)-G particles BHK-G43 cells were seeded day 1 to reach a confluence of 70-80% at day 2, where Mifepristone (10^−8^ M) was added to induce transcription of glycoprotein G. After 6 hours the medium was replaced with fresh DMEM containing 5% FBS, 50 U/mL P/S, and VSV*ΔG(luc) at MOI = 0.3. After 1 hour of incubation at 37°C BHK-G43 cells were washed three times in PBS and fresh media was added. Cells were incubated for 24 hours, after which the supernatant was centrifuged at 2000 xg for 10 min at room temperature to pellet cellular debris, and stored at −80 °C.

VSV*ΔG(luc)-SARS-2-S pseudovirus was produced by transfection with pCG1-SARS-2-S followed by transduction with VSV*ΔG(luc)-G. HEK293T cells were seeded in DMEM containing 10% FBS and 50 U/mL P/S to reach 70-80% confluence the next day. 2 μg plasmid was used per 1 × 10^6^ cells and incubated with PEI (3:1) for 30 min at room temperature. The transfection mixture was added to the cells, and incubated for 18 hours at 37 °C. Cells were washed twice with PBS, transduced with VSV*(luc)+G at MOI = 2, and incubated for 2 hours. The virus was removed by gently washing with PBS twice, and fresh DMEM containing 10% FBS and 50 U/mL P/S was added. Cell supernatant was harvested after 24 hours, centrifuged at 2000 xg for 10 min to eliminate cellular debris, and stored at −80 °C immediately. A VSV*ΔG(luc)-mock was generated simultaneously to allow subtraction of any remaining background from VSV*ΔG(luc)-G signals.

### Neutralization Assay

The SARS-CoV-2 neutralization capacity of plasma was assessed through infection of Vero76 cmyc hTMPRSS2 cells, with VSV*ΔG(luc)-SARS-2-S pseudovirus particles. Neutralization was conducted as follows: Plasma samples were thawed and heat-inactivated at 56 °C for 45 min. Subsequently, five-fold serial dilution in DMEM containing 10% FBS and 50 U/mL P/S were made. 25 μL of each plasma dilution was incubated with 50 μL VSV*ΔG(luc)-SARS-2-S at MOI = 0.01 in duplicates, for 1 hour at 37 °C, in a flat bottomed 96-well plate. Successively, 20,000 Vero76 cmyc hTMPRSS2 cells, in 50 μL DMEM containing 10% FBS and 50 U/mL P/S were added to each well, and incubated at 37 °C for 20 hours. Cells were prepared for flow cytometry by gently removing the culture media, and washing once with PBS. Cell suspensions were made by incubating each well with 75 μL Trypsin + 0.02% EDTA for 15 min at 37 °C, followed by centrifugation at 500 g for 5 min at room temperature, and re-suspension in DMEM containing 10% FBS and 50 U/mL P/S. Cells were fixed in 1% PFA for at least 15 min at 4 °C, before eGFP expression was analyzed using a Miltenyi Biotec MACSquant16 flow cytometer. The VSV*ΔG(luc)-mock eGFP background signal was subtracted from all samples.

### HLA-A2 typing and dextramer staining by flow cytometry

For HLA-A2 typing cryopreserved PBMCs were thawed, stained at room temperature for 20 min with HLA-A2 (clone BB7.2, Biolegend Cat. No. 343328) or matching isotype control (Biolegend Cat. No. 400356) and acquired on a five-laser Fortessa flow cytometer. The dextramer stains were then performed on the HLA-A2 positive samples as follows. PBMCs were incubated at room temperature for 30 min with the following SARS-CoV-2 dextramers (all from Immundex): A*0201/TLACFVLAAV-PE (Cat. No. WB3848-PE), A*0201/GMSRIGMEV-FITC (Cat. No. WB5751-FITC), A*0201/LLLDRLNQL-APC (Cat. No. WB5762-APC), A*0201/ILLNKHIDA-PE (Cat. No. WB5848-PE), A*0201/RLNEVAKNL-FITC (Cat. No. WB5750-FITC), A*0201/YLQPRTFLL-APC (Cat. No. WB5824-APC), A*0201/VLNDILSRL-PE (Cat. No. WB5823-PE), A*0201/NLNESLIDL-FITC (Cat. No. WB5850-FITC), A*0201/FIAGLIAIV-APC (Cat. No. WB5825-APC), A*0201/LLLNCLWSV-PE (Cat. No. WB3513-PE), or positive/negative control dextramers: A*0201/NLVPMVATV-PE (Cat. No. WB2132-PE, Pos. Control, CMV), A*0201/NLVPMVATV-FITC (Cat. No. WB2132-FITC, Pos. Control, CMV), A*0201/NLVPMVATV-APC (Cat. No. WB2132-APC, Pos. Control, CMV), A*0201/Neg. Control-PE (Cat. No. WB2666-PE), A*0201/Neg. Control-FITC (Cat. No. WB2666-FITC), A*0201/Neg. Control-APC (Cat. No. WB2666-APC). Cells were washed and stained with viability dye (Zombie Violet, Biolegend, Cat. No. 423114) and CD8 (Clone RPA-T8, BD, Cat. No. 563795) and acquired on a five-laser Fortessa flow cytometer.

### Data and Statistical analyses

Flow cytometry data was analyzed using FlowJo (Version 10.7.1). All data was processed and graphed in GraphPad Prism version 8.4.3. Mann-Whitney U t-test was used to compare between different groups. Spearman’s rank correlation analysis was used to access the correlation between variables as specified. Neutralization curves were plotted with three parameter non-linear fits, from which IC50 values were calculated. p ≤ 0.05 was interpreted as statistically significant. P-values are indicated as follows: n.s. = not significant,* = p ≤ 0.05, ** = p < 0.01, *** = p < 0.001, **** = p < 0.0001.

## Supporting information

Supportive information

## Acknowledgements

We would like to thank all the individuals in the study for the kind donation of both their time and biological material. Thank you to MesoScale Discovery, for providing the reagents and materials to enable this study. Thank you to Markus Hoffmann and Stefan Pöhlmann for the kind gift of BHK-G43 and Vero76 cmyc hTMPRSS2 cells and the pCGI-SARS-2-spike plasmid. Figure 5A was created with the help of BioRender.com. We thank Lene Svinth Jønke for her immense assistance in the laboratory during patient material collection. Thank you, to the entire staff of the Department of Infectious Diseases, for their feedback and scientific discussions.

## Author contributions

SFN, LKV, MT, MHS, OSS, LØ contributed to study design, data collection, data analysis, data interpretation, literature search, and the writing of this report. IMJ, RO, GSF, MHP, CE, AH, and RH contributed to experiments, data analysis, and data interpretation. JFH, JDG and LKV contributed to individual recruitment, data collection and clinical management. The final version of this paper was reviewed and approved by all authors.

