## Supplementary material for "SARS-CoV-2 elicits robust adaptive immune responses regardless of disease severity": Supportive information

**S1 Appendix**

**Overview: Page no.**

1. **Figure A: Time from recovery to inclusion correlates with antibody volume.**
2. **Figure B: Full gating strategy for CD8+ T cell dextramer staining.**
3. **Figure C: Distribution of non-CD8+ T-cell responders IC50 values within the cohort:**

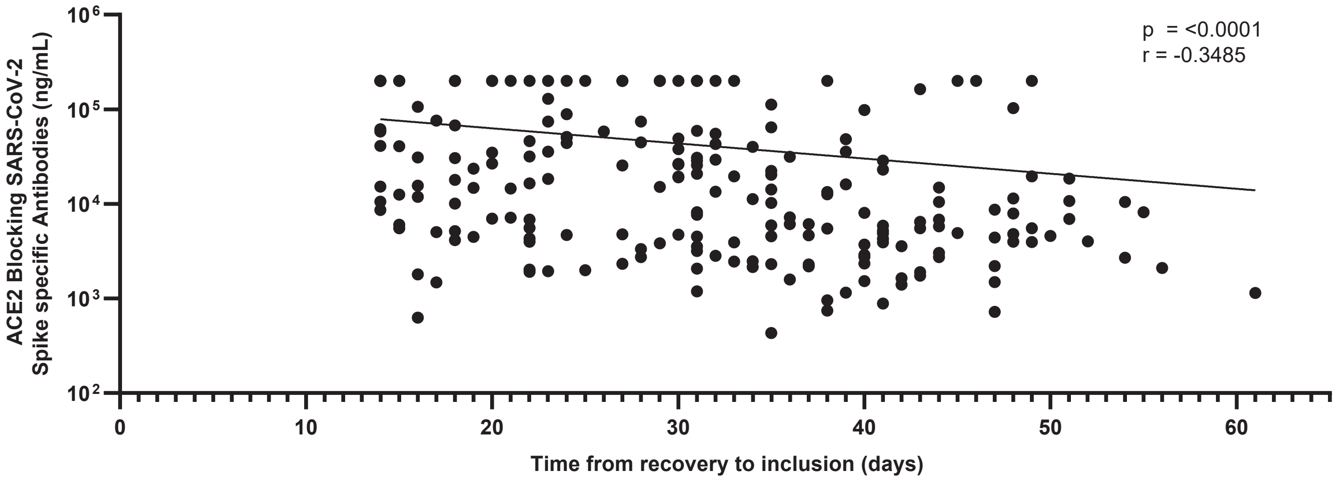


**Appendix Figure A: Time from recovery to inclusion correlates with antibody volume.** The time from recovery to inclusion in days (x) plotted against ACE2 Blocking SARS-CoV-2 Spike specific antibodies in ng/mL (y). Correlation by Spearman's rank coefficient, $p < 0.0001$. n = 203

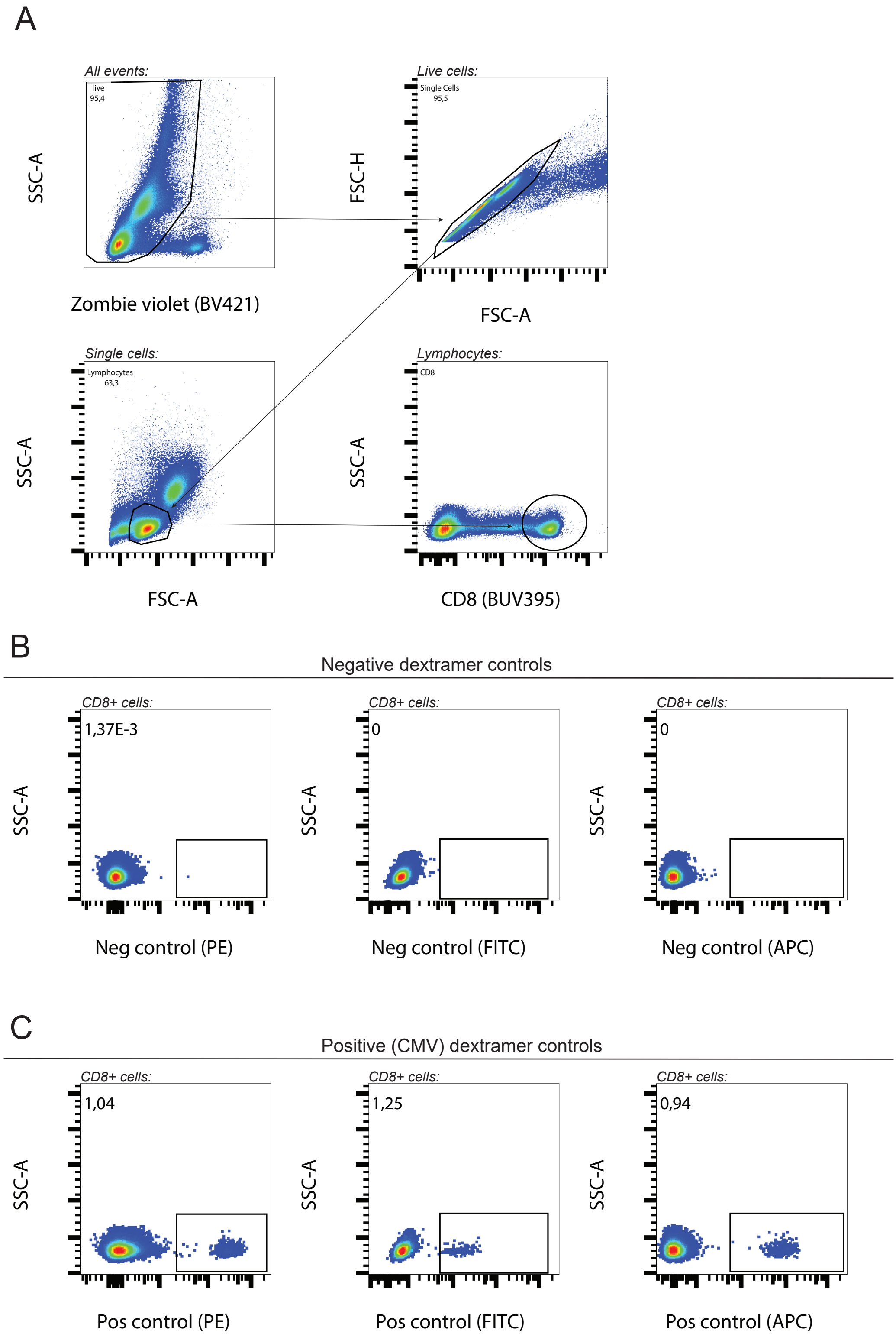


**Appendix Figure B: Full gating strategy for CD8+ T cell dextramer staining. A)** Gating strategy for the isolation of first live cell events, single cells, lymphocytes and finally CD8+ T cells. **B)** Negative control dextramer gates for PE, FITC and APC fluorophores. **C)** Positive control dextramer gates for PE, FITC and APC fluorophores.

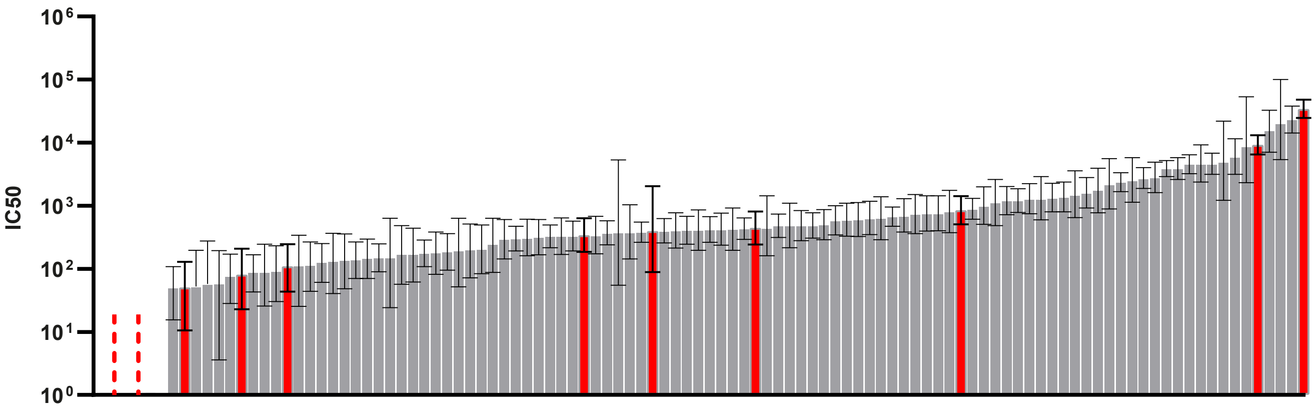


**Appendix Figure C: Distribution of non-CD8+ T-cell responders IC50 values within the cohort:** Listed IC50 values for all HLA-A2+ individuals, ranked from lowest to highest. Individuals without a detectable CD8+ T-cell response to any of the applied dextramer epitopes are shown in red. Six HLA-A2+ individuals were unable to neutralize pseudovirus 100 %, and have no IC50 values assigned. Two of them showed no detectable CD8+ T-cell response, and are schematically illustrated by red dotted lines.
